# STK25 is an IRF5 kinase that promotes TLR7/8-mediated inflammation

**DOI:** 10.1101/2023.09.26.559637

**Authors:** Matthew R. Rice, Bharati Matta, Loretta Wang, Surya Indukuri, Betsy J. Barnes

## Abstract

Toll-like receptors (TLRs) represent a subset of pattern-recognition receptors (PRRs) employed by the innate immune system to detect pathogen-associated molecular patterns (PAMPs) and initiate the response to invading microbes. The transcription factor interferon regulatory factor 5 (IRF5) functions as an important mediator of the inflammatory response downstream of MyD88-dependent TLR activation. While the dysregulation of IRF5 activity has been implicated in the development of several autoimmune diseases including systemic lupus erythematosus (SLE) and rheumatoid arthritis, the factors that modulate TLR-induced IRF5 post-translational modifications (PTMs) are poorly understood. Therefore, the focus of this study was to identify and characterize the role(s) of novel kinases in the regulation of TLR7/8 signaling. We performed a kinome-wide siRNA screen in human THP-1 monocytic cells to identify mediators of TLR7/8-induced TNF-α and IL-6 production. We identified serine/threonine protein kinase 25 (STK25) as a positive regulator of proinflammatory cytokine release in response to TLR7/8 activation in human primary myeloid cells. We determined that STK25 phosphorylates IRF5 *in vitro* via multiple biochemical assays. Phosphopeptide mapping by mass spectrometry revealed that STK25 phosphorylates IRF5 at a highly conserved residue, Thr265, that leads to the transcriptional activation of IRF5 in HEK293T cells. We determined that STK25 undergoes autophosphorylation in response to a variety of TLR triggers in multiple immune cell types. We demonstrated that R848-induced IRF5 nuclear translocation and proinflammatory cytokine production was significantly attenuated in immune cells from *Stk25*-deficient mice compared to wild-type. Finally, we determined that STK25 autophosphorylation is increased at steady-state in peripheral blood mononuclear cells (PBMCs) from SLE donors compared to healthy controls. Thus, our findings implicate STK25 as an important regulator of TLR7/8 signaling through the modulation of IRF5 activation.

**Significance Statement:** The transcription factor IRF5 functions as a master regulator of innate and adaptive immunity. While the hyperactivation of IRF5 has been implicated in the pathogenesis of systemic lupus erythematosus (SLE), the mechanisms leading to the modulation of IRF5 activity are incompletely understood. Here, we conducted a screen of the human kinome to identify IRF5 kinases that function as positive regulators of TLR-induced inflammation. We demonstrate that STK25 directly phosphorylates IRF5 to drive proinflammatory cytokine responses downstream of TLR activation in both human and murine primary immune cells. Altogether, our findings implicate STK25 as a potential therapeutic target for the management of IRF5-mediated immunological disorders.

## Introduction

The innate immune system utilizes pattern-recognition receptors (PRRs) to sense the presence of invading pathogens. Toll-like receptors (TLRs) are an important class of PRRs that recognize conserved microbial elements known as pathogen-associated molecular patterns (PAMPs) and host-derived signals from injured cells called damage-associated molecular patterns (DAMPs). TLR engagement facilitates the activation of signaling pathways that culminate in the induction of proinflammatory cytokines and type I interferons (IFNs) (1). The TLR-induced production of proinflammatory mediators promotes microbial clearance and primes the secondary pathogen-specific adaptive immune response (1). The transcription factor interferon regulatory factor 5 (IRF5) functions as a critical modulator of the inflammatory response downstream of MyD88-dependent TLR activation. The activation of IRF5 involves a series of post-translational modifications (PTMs) that include TRAF6-mediated ubiquitination and phosphorylation by IKKβ (2, 3, 4). In turn, a network of negative regulatory factors, including Lyn, IRF4, and IKKα, function to restrain the activation of IRF5 downstream of TLR ligation (5, 6, 7). Genetic polymorphisms within or near the *IRF5* locus have been associated with IRF5 hyperactivation and the dysregulation of IRF5 activity has been implicated in the pathogenesis of systemic lupus erythematosus (SLE) (8). Given that dysregulated functions of TLRs have also been implicated in SLE, the identification of kinases that regulate the TLR-mediated activation of IRF5 may lead to the development of therapeutic agents for the treatment of patients with autoimmune and inflammatory conditions.

Serine/threonine protein kinase 25 (STK25) is a member of the germinal center kinase (GCK)-III subfamily of the mammalian sterile 20-like (MST) kinase family. STK25 has been implicated in the regulation of metabolic homeostasis, neuronal polarization, cell migration, Golgi organization, apoptosis, and tumorigenesis (9, 10, 11, 12, 13). However, a role for STK25 in the modulation of TLR signaling has yet to be defined.

In this study, we screened for putative IRF5 kinases involved in the regulation of TLR7/8-mediated proinflammatory cytokine release. Of the candidate kinases identified, only STK25 directly phosphorylated IRF5 *in vitro*. STK25 promoted IRF5 transcriptional activation in cells via phosphorylation of Thr265. In addition, signaling through TLR7/8 induced the expression and activation of STK25. Furthermore, STK25 positively regulated TLR7/8-induced IRF5 nuclear translocation and proinflammatory factor production in murine primary immune cells. Finally, peripheral blood mononuclear cells (PBMCs) from patients with SLE demonstrated elevated expression of autophosphorylated STK25 as compared to healthy controls. Altogether, our findings support a role for STK25 in the regulation of TLR signaling and thereby implicate STK25 as a potential therapeutic target for the treatment of IRF5-mediated immunological disorders.

## Results

### Identification of positive regulators of TLR7/8-induced proinflammatory cytokine production

To identify candidate kinases involved in the regulation of IRF5 downstream of TLR7/8-mediated signaling, we conducted a kinome-wide siRNA screen of the THP-1 human monocytic cell line (Figure 1A). Each kinase was targeted by a single siRNA molecule at a time, for a total of 3 distinct siRNA constructs per target. The cells were stimulated with R848, a TLR7/8 agonist, 48 h after siRNA knockdown, and culture supernatants were harvested 24 h post-stimulation. Alterations in the R848-induced production of TNF-α and IL-6 following siRNA knockdown were determined by AlphaLISA immunoassay. We used strictly standardized mean difference (SSMD) scores to compare the effect of each siRNA molecule on R848-induced proinflammatory cytokine production. An SSMD score of -1 for a given siRNA molecule was considered a positive hit and genes with 2 or more hits were selected for follow up analysis. The normal distribution of the SSMD scores indicated that most kinases were not involved in the R848-induced production of proinflammatory mediators (Figure S1A). As expected, the knockdown of IRF5 with 2 independent siRNA constructs led to a significant reduction in the production of TNF-α and IL-6 in response to R848 (Figure S1B). The identification of well-established TLR signaling components, interleukin-1 receptor-associated kinase 1 (IRAK1) and IRAK2 as positive regulators of the inflammatory response to R848, provided validation for the screen (Figure S1B). The 20 kinases that exhibited the greatest reduction in R848-induced proinflammatory cytokine production following siRNA knockdown are ranked by SSMD score in Figure 1A. While many of the hits represent known modulators of TLR signaling, serine/threonine protein kinase 16 (STK16), STK25, mitogen-activated protein kinase kinase kinase 19 (MAP3K19), and sphingosine kinase 2 (SPHK2), were selected for validation in follow up studies as they had never been associated with TLR7/8-induced cytokine production. Given that siRNA knockdown of STK16, STK25, MAP3K19, and SPHK2 in THP-1 cells reduced the R848-induced production of proinflammatory cytokines, we sought to confirm the relevance of these findings in human primary myeloid cells. We isolated monocytes from the peripheral blood of healthy donors and generated monocyte-derived dendritic cells (MDDCs) via incubation with GM-CSF and IL-4. The cells were subjected to targeted siRNA knockdown for 48 h and then stimulated with R848 for 24 h. The culture supernatants were harvested and the production of TNF-α and IL-6 was quantified via AlphaLISA immunoassay. Interestingly, knockdown of STK25 in human primary monocytes and MDDCs led to a substantial reduction in the R848-mediated production of TNF-α and IL-6 (Figure 1B-E). Notably, knockdown of STK16, MAP3K19, and SPHK2 led to a similar attenuation of R848-induced TNF-α and IL-6 release in MDDCs, with varying effects in human primary monocytes (Figure 1B-E).

**Figure 1.**
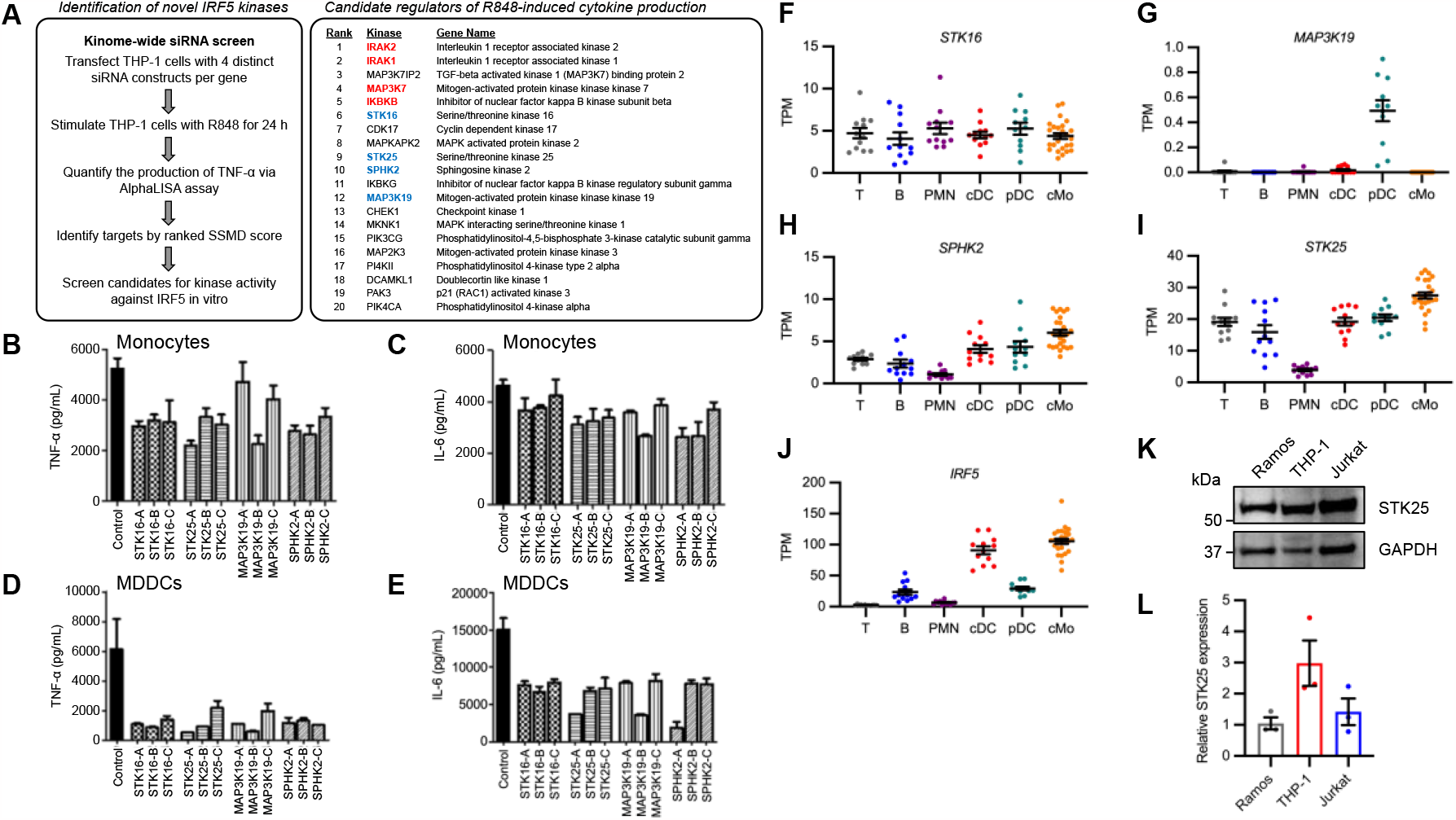
Identification of kinases involved in the regulation of R848-induced inflammation. **A**, Overview of the siRNA screen conducted in THP-1 cells to identify kinases that modulate R848-mediated proinflammatory cytokine release. The 20 kinases that exhibited the most significant reduction in R848-induced TNF-α and IL-6 production following siRNA knockdown are listed by ranked SSMD score. Previously characterized regulators of TLR signaling are highlighted in red. Targets displayed in blue have never been associated with innate immune signaling. **B-C**, Quantification of TNF-α (**B**) and IL-6 **(C**) production in culture supernatants of human primary monocytes stimulated with R848 for 24 h following siRNA knockdown of *STK16, STK25, MAP3K19*, or *SPHK2* (*N* = 3 technical replicates with 3 distinct siRNA constructs per target). Data represent mean ± SEM. **D-E**, Quantification of TNF-α (**D**) and IL-6 **(E**) production in culture supernatants of human primary monocyte-derived dendritic cells (MDDCs) stimulated with R848 for 24 h following siRNA knockdown of *STK16, STK25, MAP3K19*, or *SPHK2* (*N* = 3 technical replicates with 3 distinct siRNA constructs per target). **F-J**, Relative mRNA expression of *STK16* (**F**), *MAP3K19* (**G**), *SPHK2* (**H**), *STK25* (**I**), and *IRF5* (**J**) in sorted peripheral blood leukocytes from healthy control donors (*N* = 11-26). Normalized transcript per million (TPM) values were obtained from GSE149050. T, T cells; B, B cells; PMN, neutrophils; cDC, conventional dendritic cells; pDC, plasmacytoid dendritic cells; cMo, classical monocytes. **K**, Immunoblot analysis of STK25 protein expression in untreated human cancer cell lines. Blots were probed with antibodies against STK25 and GAPDH. **L**, Densitometric analysis of STK25 protein levels after normalization to the expression of GAPDH (*N* = 3 biological replicates). Data represent mean ± SEM.

### *STK25* is highly expressed in multiple immune cell types

Since STK16, STK25, MAP3K19, and SPHK2 have never been implicated in the regulation of TLR7/8-mediated cytokine production, we utilized a published RNA-seq dataset to analyze the gene expression levels of each kinase in sorted peripheral blood cell populations from healthy human donors (14). The expression levels of *STK16, MAP3K19*, and *SPHK2* were particularly low in most of the examined immune cell populations with a mean TPM < 8 (Figure 1F-H). However, *STK25* was expressed in nearly all immune cell populations with a mean TPM > 15, except for neutrophils (Figure 1I). We found that *STK25* was highly expressed in classical monocytes, a cell type that exhibits the highest expression of *IRF5* (Figure 1I-J). We next examined STK25 protein expression in a spectrum of human hematologic malignancies (B cell lymphoma, T cell leukemia, and monocytic leukemia) by immunoblot analysis and determined that STK25 was expressed in all cell types, with the highest expression in THP-1 monocytes (Figure 1K-L). In summary, STK25 was broadly expressed in human immune cells, suggesting a previously uncharacterized role for STK25 in the regulation of immune cell function. Furthermore, STK25 was highly expressed in human monocytes, a cell type that is critical to the innate immune response to pathogens.

### STK25 induces IRF5 transcriptional activation via phosphorylation at Thr265

Given the importance of IRF5 PTMs in the modulation of TLR-induced transcriptional activation, we evaluated the ability of each target kinase to phosphorylate a biotinylated C-terminal construct (residues 222-467) of IRF5 via an *in vitro* scintillation proximity assay. The phosphorylation of IRF5 by IKKβ and IRAK1 provided validation for the assay (Figure S2A). Of the candidate kinases, only STK25 possessed the ability to phosphorylate the truncated form of IRF5 (Figure 2A). Additionally, we found that IKKβ and STK25 could phosphorylate full-length IRF5 in a dose-dependent manner *in vitro* via a luminescent kinase assay system (Figure 2B-E). Finally, we evaluated the kinetics of STK25-mediated and IKKβ-mediated IRF5 phosphorylation *in vitro* by Phos-tag immunoblot analysis. Interestingly, while the kinetics of IRF5 phosphorylation by STK25 and IKKβ were similar, with detectable IRF5 phosphorylation within 30 min, and substantial phosphorylation by 60 min, the patterns of modification appeared different suggesting phosphorylation at distinct sites (Figure 2F). Since STK25 functions as a serine/threonine kinase, we sought to determine which IRF5 residues are specifically modified by STK25. Immunoblot analysis of a 60 min kinase reaction with STK25 and IRF5 revealed that STK25 phosphorylated IRF5 at threonine (Thr) residues (Figure 2G). To identify the specific IRF5 residues that are phosphorylated by STK25, we employed mass spectrometry. We found that STK25 phosphorylated IRF5 at multiple residues, including Thr183, Thr265, and Thr314 (Figure 2H and S3A). Since these IRF5 residues have not been previously implicated in the regulation of IRF5 activity, we aligned IRF5 protein sequences from characterized human isoforms and multiple species to evaluate conservation. Interestingly, Thr265 is completely conserved across all human isoforms and queried species, while Thr183 is conserved across most human isoforms and substituted with an Ala residue in *M. musculus* (Figure 2I-J and S3B-C). The evolutionary conservation of Thr265 suggests that this residue may be important for IRF5 function. Therefore, STK25 is a putative IRF5 kinase that phosphorylates IRF5 at a highly conserved residue, Thr265. While the phosphorylation of serine residues in the C-terminal region of IRF5 has been demonstrated to regulate IRF5 activity, the functional significance of Thr phosphorylation is poorly understood (3, 4). As such, we utilized an interferon-stimulated response element (ISRE) firefly luciferase (ISRE-Luc) reporter system to evaluate STK25-mediated IRF5 activation in HEK293T cells. We observed basal ISRE-Luc activity with the expression of wild-type IRF5 (WT-IRF5) (Figure 2K). However, the co-expression of IRF5 and STK25 led to a significant increase in ISRE-Luc activity, indicating that STK25 promoted IRF5 transcriptional activity (Figure 2K). We performed site-directed mutagenesis to generate an alanine-substituted mutant form of IRF5 at residue 265 (IRF5-T265A) that would be resistant to STK25-mediated phosphorylation. Interestingly, the IRF5-T265A mutant exhibited reduced basal ISRE-Luc activity compared to WT-IRF5 (Figure 2K). While the co-expression of IRF5-T265A and STK25 led to a slight increase in ISRE-Luc activity compared to IRF5-T265A alone, the level of activation was not significantly different from IRF5-T265A or WT-IRF5 alone (Figure 2K). Importantly, the observed differences in the activation of WT-IRF5 and IRF5-T265A were not due to altered expression of the IRF5-T265A mutant (Figure 2L). Altogether, these findings support a role for Thr265 in STK25-mediated IRF5 transcriptional activation in cells.

**Figure 2.**
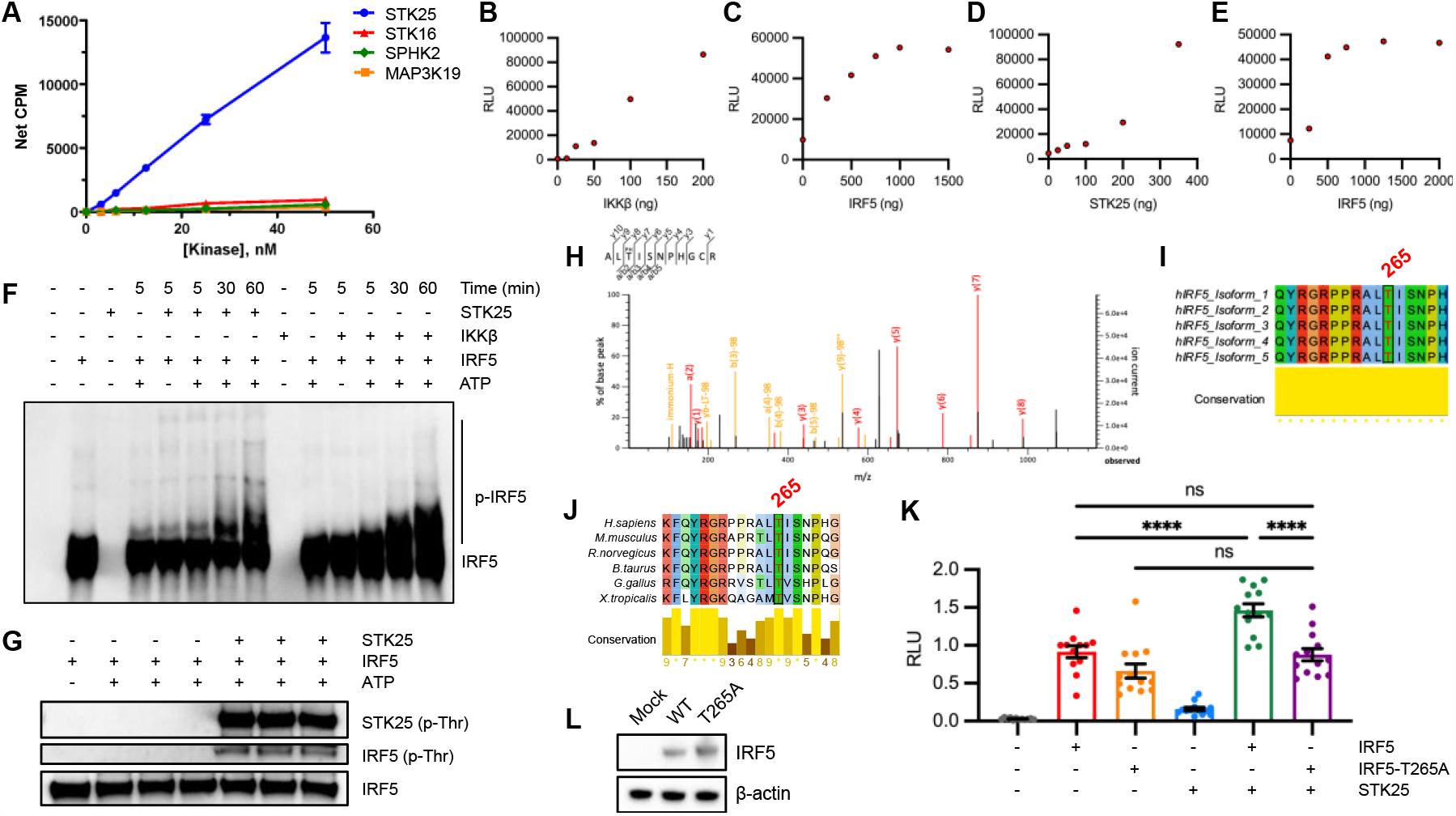
STK25 induces IRF5 transcriptional activation via phosphorylation at Thr265. **A**, Phosphorylation of a biotinylated C-terminal construct of IRF5 (residues 222-467) by candidate kinases in an *in vitro* scintillation proximity assay (*N* = 4 biological replicates). **B-E**, Phosphorylation of full-length IRF5 by IKKβ (**B-C**) or STK25 (**D-E**) in an *in vitro* luminescent kinase assay. **B**, Dose-dependent phosphorylation of IRF5 (500 ng) by IKKβ. **C**, Dose-dependent phosphorylation of IRF5 by a fixed amount of IKKβ (100 ng). **D**, Dose-dependent phosphorylation of IRF5 (500 ng) by STK25. **E**, Dose-dependent phosphorylation of IRF5 by a fixed amount of STK25 (350 ng). **F**, Phostag immunoblot analysis of the kinetics of STK25- and IKKβ-mediated phosphorylation of full-length IRF5 *in vitro*. Blots were probed with an antibody against total IRF5. p-IRF5, phosphorylated IRF5. **G**, Immunoblot analysis of IRF5 phosphorylation at Thr residues following incubation with STK25 in an *in vitro* kinase assay for 1 h at RT. Blots were probed with antibodies against total p-Thr, p-STK25 (T174), and IRF5. Representative of 3 independent experiments. p-Thr, phosphorylated-Thr. **H**, Mass spectrometry-based identification of Thr265 as an STK25-dependent IRF5 phosphorylation site. *In vitro* kinase reactions with IRF5 and STK25 were incubated for 1 h at RT, subjected to SDS-PAGE, and the gel was stained with Coomassie blue. Protein bands were excised, destained, and digested with trypsin. Representative of 3 independent experiments. **I**, Conservation of Thr265 across multiple human isoforms of IRF5. **J**, Conservation of Thr265 in IRF5 protein sequences from multiple species. **K**, HEK293T cells were co-transfected with IRF5-FLAG, IRF5-T265A-FLAG, or STK25 in addition to an interferon-stimulated response element (ISRE) firefly luciferase (ISRE-Luc) reporter and a cytomegalovirus (CMV) *Renilla* luciferase (CMV-RL) internal control. HEK293T cells were also co-transfected with STK25 in combination with IRF5-FLAG or IRF5-T265A-FLAG, along with an ISRE-Luc reporter and CMV-RL internal control. The firefly luciferase signal was normalized to the *Renilla* luciferase signal to determine the relative luciferase units (RLU). For each experiment, the relative ratio for each sample was normalized to a sample with IRF5-FLAG, ISRE-Luc, and CMV-RL (*N* = 12-13 transfections from 3 independent experiments). \*\*\*\**P* < 0.0001. **L**, Immunoblot analysis of HEK293T cells co-transfected with IRF5-FLAG or IRF5-T265A-FLAG. Blots were probed with antibodies against IRF5 and β-actin. Data represent mean ± SEM.

### STK25 responds to TLR7/8 activation in THP-1 cells and regulates TLR-induced proinflammatory cytokine production in murine primary immune cells

All members of the GCK-III subfamily require the phosphorylation of a conserved Thr residue within the activation T-loop for full kinase activity (15). The phosphorylation of STK25 at Thr174 (T174) is hypothesized to be the result of a *trans*-autophosphorylation event that involves STK25 homodimerization (15). As such, we examined whether TLR7/8 activation induces STK25 autophosphorylation at Thr174 in THP-1 cells by immunoblot analysis. Despite the detection of basal levels of phospho-STK25 (T174) (p-STK25) in untreated cells, we observed an increase in p-STK25 expression at 30 min post-stimulation with R848 (Figure 3A-B). Importantly, R848-induced STK25 autophosphorylation occurred prior to IRF5 nuclear translocation, which has been previously detected at 2 h post-stimulation in THP-1 cells, amongst other cell types (16, 17, 18). Interestingly, STK25 activation was not restricted to TLR7/8 signaling, as treatment with LPS, a TLR4 ligand, induced STK25 autophosphorylation at 6 h post-stimulation, and treatment of B cells with CpG-B, a TLR9 ligand, induced autophosphorylation within 30 min (Figure S4A-D). These data suggest that STK25 becomes activated downstream of multiple MyD88-dependent TLRs and may regulate additional intracellular signaling cascades in other immune cell types. In addition, we found that STK25 protein expression was significantly upregulated in THP-1 cells at 24 h post-stimulation with R848 (Figure 3C-D). Together, these findings indicate that STK25 is modulated at two levels (phosphorylation and protein expression) downstream of TLR7/8 signaling. To further characterize the role of STK25 in TLR-induced IRF5 activation in immune cells, we generated *Stk25*-deficient (*Stk25*^*-/-*^) mice by Cre-mediated excision of exons 4 and 5 as previously described (10). Deletion of *Stk25* was confirmed at the protein level by immunoblot analysis of *Stk25*^*+/+*^ (WT) and *Stk25*^*-/-*^ (KO) splenocytes (Figure 3E). Given the ability of STK25 to modulate R848-induced proinflammatory cytokine production in siRNA knockdown studies in human primary myeloid cells, we sought to further validate these findings via parallel experiments in immune cells from WT and KO mice. We isolated peripheral blood cells from WT and KO mice and evaluated IRF5 nuclear translocation at 2 h post-stimulation with R848 or CpG-B by imaging flow cytometry. We observed a significant reduction in R848-induced IRF5 nuclear translocation in KO peripheral blood CD11b^+^ monocytes and B220^+^ B cells compared to WT (Figure 3F-G). We also found a significant reduction in CpG-B-induced IRF5 nuclear translocation in KO peripheral blood B220^+^ B cells relative to WT (Figure 3G). By intracellular flow cytometry, we observed a substantial reduction in the frequencies of IL-6^+^CD11b^+^ and TNF-α^+^CD11b^+^ splenocytes from KO mice compared to WT in response to an 18 h stimulation with R848 (Figure 3H-I). To evaluate the role of TLR specificity in STK25-mediated inflammatory responses, WT and KO splenocytes were stimulated with R848, CpG-B, or LPS for 24 h *in vitro* and the production of IL-6 in culture supernatants was quantified via ELISA. Importantly, both R848- and CpG-B-induced production of IL-6 was significantly attenuated in KO splenocytes compared to WT (Figure 3J-K). However, LPS-induced production of IL-6, although reduced in KO splenocytes, was not found to be statistically different than WT splenocytes (Figure 3L). These findings supplement the results of our initial siRNA-based studies in human primary myeloid cells and provide additional insight into the conservation of STK25 function across multiple species and TLR signaling pathways.

**Figure 3.**
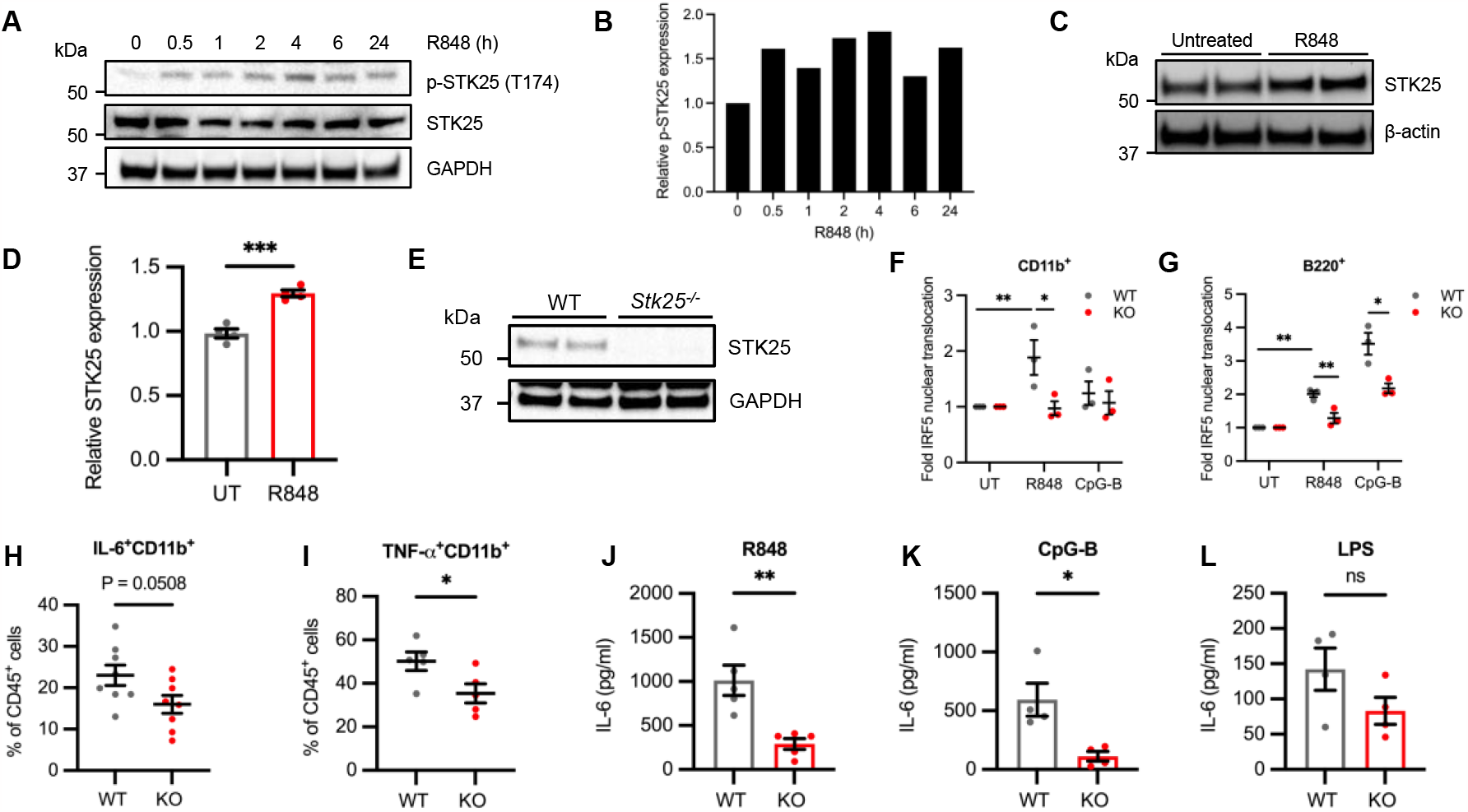
STK25 responds to TLR7/8 activation in THP-1 cells and regulates TLR-induced proinflammatory cytokine production in murine primary immune cells. **A** Immunoblot analysis of STK25 autophosphorylation at Thr174 in THP-1 cells stimulated with R848 for 0.5, 1, 2, 4, 6, or 24 h. Representative of 2 independent experiments. Blots were probed with antibodies against p-STK25 (T174), STK25, and GAPDH. **B**, Representative densitometric analysis of p-STK25 (T174) protein levels after normalization to total STK25 protein levels and the expression of GAPDH. **C**, Immunoblot analysis of STK25 expression in THP-1 cells stimulated with R848 for 24 h. **D**, Densitometric analysis of STK25 protein levels after normalization to the expression of β-actin (*N* = 4 biological replicates). UT, untreated. **E**, Immunoblot analysis of STK25 protein expression in *Stk25*^*+/+*^ (WT) and *Stk25*^*-/-*^ (KO) splenocytes. Blots were probed with antibodies against STK25 and GAPDH (*N* = 2 biological replicates per genotype). **F-G**, Imaging flow cytometry analysis of IRF5 nuclear translocation in peripheral blood CD11b^+^ monocytes (**F**) and B220^+^ B cells (**G**) from WT and KO mice following stimulation with R848 or CpG-B for 2 h (*N* = 3 biological replicates per genotype). **H-I**, Flow cytometry analysis of the frequencies of IL-6^+^CD11b^+^ (**H**) and TNF-α^+^CD11b^+^ (**I**) splenocytes from WT and KO mice following stimulation with R848 for 18 h (*N* = 5-8 biological replicates per genotype). **J-L**, Quantification of IL-6 production in culture supernatants of WT and KO splenocytes stimulated with R848 (**J**), CpG-B (**K**), or LPS (**L**) for 24 h (*N* = 4-5 biological replicates per genotype). \**P* < 0.05, \*\**P* < 0.01, \*\*\**P* < 0.001. Data represent mean ± SEM.

### Basal autophosphorylation of STK25 at Thr174 is increased in PBMCs from patients with SLE

While the activation of IRF5 plays a vital role in both the innate and adaptive arms of the immune system, IRF5 hyperactivation contributes to the development of autoimmunity (19, 20). In multiple murine models of SLE, inhibition of IRF5 attenuates disease activity, and thus IRF5 has emerged as an important therapeutic target (21, 22, 23). Given the ability of STK25 to induce the activation of IRF5, we sought to determine whether STK25 is autophosphorylated in immune cells from SLE patients at steady-state. To begin to address this hypothesis, we obtained PBMCs from a small cohort of healthy controls and patients with SLE and measured total STK25 and p-STK25 expression by immunoblot analysis. Interestingly, we observed a substantial increase in p-STK25 expression in SLE PBMCs compared to healthy controls, despite similar expression levels of total STK25, suggesting that STK25 is hyperactivated in SLE PBMCs (Figure 4A-C). Altogether, these findings implicate STK25 as a potential driver of IRF5 hyperactivation in SLE (Figure 4D).

**Figure 4.**
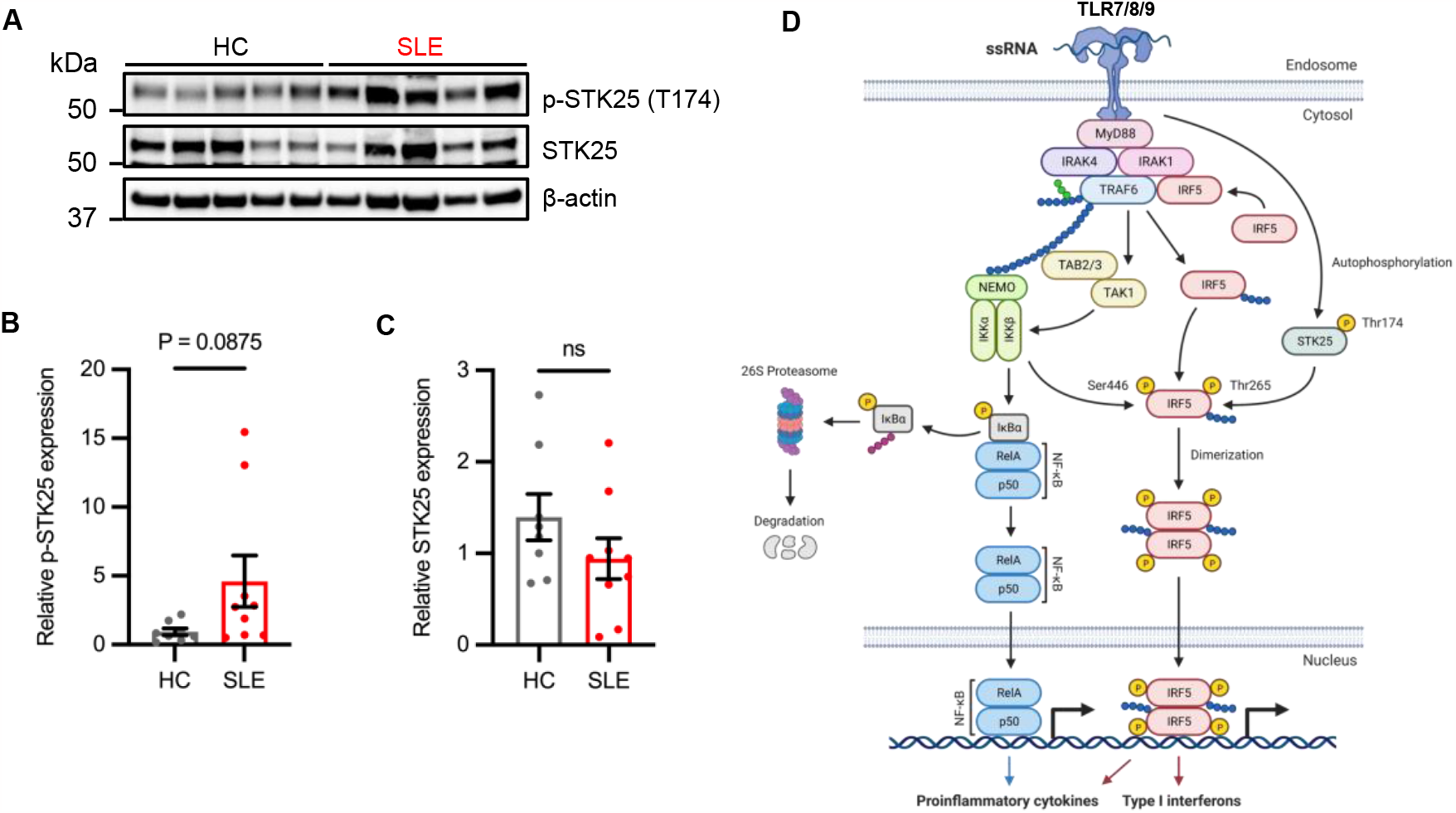
Basal autophosphorylation of STK25 at Thr174 is increased in PBMCs from patients with SLE. **A**, Immunoblot analysis of STK25 autophosphorylation at Thr174 in PBMCs from healthy donors and patients with SLE. (*N* = 8-9 samples per condition). Blots were probed with antibodies against p-STK25 (T174), STK25, and β-actin. **B**, Densitometric analysis of total STK25 protein levels after normalization to the expression of β-actin. **C**, Densitometric analysis of p-STK25 (T174) protein levels after normalization to total STK25 protein levels and the expression of β-actin. Data represent mean ± SEM. **D**, Proposed model for STK25 as an IRF5 kinase downstream of TLR7/8/9 signaling. Created with BioRender.com.

## Discussion

In our efforts to characterize critical regulators of IRF5 downstream of MyD88-dependent TLR activation, we identified 4 new kinases - STK16, MAP3K19, SPHK2, and STK25 that when knocked down, led to a reduction in the R848-induced production of TNF-α and IL-6 in THP-1 monocytes, human primary monocytes, and MDDCs (Figure 1A-E). Interestingly, the degree of inhibition of R848-induced proinflammatory cytokine production was cell type-specific, with a more profound effect observed in MDDCs for all candidate kinases (Figure 1D-E). While the reason for the observed differences in the ability of these kinases to regulate proinflammatory cytokine responses in human primary monocytes and MDDCs is currently unknown, the cell type-specific expression pattern of each kinase may be involved. Interestingly, gene expression profiling of peripheral leukocytes from healthy donors indicated that *STK25* was highly expressed in multiple cell types, especially classical monocytes (Figure 1I). In humans, classical monocytes are a subset of monocytes that utilize TLRs to initiate the inflammatory response to pathogens (24). The expression of *IRF5* was also elevated in classical monocytes and the aberrant activation of IRF5 in monocytes has been implicated in the development of several autoimmune conditions, including SLE (8). Furthermore, the examination of STK25 protein expression in human cell lines supported the gene expression profiles in human primary cells, as STK25 was highly expressed in THP-1 cells (Figure 1K-L). These findings are in agreement with previous reports of STK25 expression in non-hematopoietic cell types, in that STK25 was ubiquitously expressed in most tissues (25). Altogether, we have demonstrated that STK25 is expressed in multiple immune cell subsets, with the highest levels of expression in TLR-responsive monocytes.

Since the activation of TLR7/8 by R848 results in the IKKβ-dependent activation of both NF-κB and IRF5, we examined whether the regulation of proinflammatory cytokine production by STK16, MAP3K19, SPHK2, and STK25 was a direct result of their ability to regulate IRF5 activation. We determined that only STK25 could phosphorylate IRF5 in a series of biochemical assays (Figure 2A-E). Thus, the ability of STK16, MAP3K19, and SPHK2 to mediate TLR-induced proinflammatory cytokine production may be due to their regulation of NF-κB activity as opposed to IRF5. Alternatively, these kinases could potentially phosphorylate the N-terminal region of IRF5 between residues 1-221, as the screening assay was performed with a truncated form of IRF5 that only contained residues 222-467. While the phosphorylation of IRF5 at Ser residues within the SRR is important for the IKKβ-mediated activation of IRF5, the phosphorylation of IRF5 at Tyr172 by PYK2 has recently been shown to promote IRF5 transcriptional activity (26). Interestingly, Tyr172 is absent from isoform 1 of IRF5, so the isoform-specific regulation of IRF5 by putative kinases should also be considered. Therefore, the ability of STK16, MAP3K19, and SPHK2 to phosphorylate full-length IRF5 should be further evaluated in future studies.

To further our understanding of the regulation of IRF5 by STK25, we mapped STK25 target residues by mass spectrometry. We found that STK25 phosphorylated IRF5 at several threonine residues, including Thr183, Thr265, and Thr314 (Figure 2H and S3A). As a serine/threonine kinase, STK25 has been shown to catalyze the phosphorylation of serine and threonine residues, with a preference for serine residues (27). Despite the regulation of multiple cellular processes by STK25, only a few direct targets of STK25 have been identified. In addition to autophosphorylation at Thr174, STK25 phosphorylates CCM3 at Ser39 and Thr43 (28). Moreover, STK25 phosphorylates 14-3-3ζ, Salvador homolog-1 (SAV1), large tumor suppressor 2 (LATS2), and cAMP-dependent protein kinase type I-alpha regulatory subunit (PRKAR1A) at Ser58, Thr26, Ser872, and Ser77/Ser83, respectively (11, 29, 30, 31). Given the importance of Ser residues within the SRR of IRF5 that regulate its activity, it was intriguing that STK25 preferentially targeted Thr residues in IRF5 over Ser residues (32, 33). Importantly, the STK25-dependent phosphorylation of IRF5 at Thr265 was robustly detected in all experimental replicates with conclusive localization (Figure 2H and S3A). To investigate the functional significance of phosphorylation at Thr265, we examined IRF5 protein sequences from several species. We found that Thr265 was conserved in all analyzed organisms and in all characterized human isoforms of IRF5 (Figure 2I-J). In contrast, Thr183 is replaced with an Ala residue in murine IRF5 and is not completely conserved across human isoforms of IRF5 (Figure S3B-C). While the phosphorylation of IRF5 represents an important step in IRF5 activation, the TLR-mediated expression of proinflammatory cytokines requires the induction of transcriptional activity (16). To examine whether STK25 modulates the ability of IRF5 to bind to an ISRE reporter, we co-expressed STK25 and IRF5 in HEK293T cells. We found that in the absence of stimulation, STK25 promoted the transcriptional activity of IRF5 (Figure 2K). To evaluate the importance of Thr265 in IRF5 activation, we mutated Thr265 to an Ala residue, and co-expressed the IRF5-T265A mutant and STK25. Importantly, the mutation of Thr265 to Ala prevented the induction of IRF5 transcriptional activity by STK25, and this was not due to aberrant expression of the IRF5-T265A mutant as we found that the IRF5-T265A mutant was normally expressed in HEK293T cells (Figure 2K-L). To our knowledge, this is the first report of kinase-mediated transcriptional activation of IRF5 via regulation of Thr265.

Although we have characterized the STK25-mediated activation of IRF5 downstream of TLR engagement, the upstream mechanisms that govern the activation of STK25 have never been examined. Autophosphorylation of STK25 at Thr174 represents an important step in kinase activation (15). Thus, we examined whether TLR activation leads to the phosphorylation of STK25 in THP-1 cells. Interestingly, we found that STK25 autophosphorylation was increased post-stimulation with R848, LPS, and CpG-B (Figure 3A-B and Figure S4A-D). These data suggest that STK25 becomes activated downstream of multiple MyD88-dependent TLRs and may regulate additional intracellular signaling cascades in other immune cell types.

STK25 was originally identified as an oxidant stress response kinase due to its ability to become activated by reactive oxygen intermediates in Ramos B cells (34). Subsequent studies in rodent L6 myoblasts and human medulloblastoma D283MED-TrkA (MB-TrkA) cells revealed that STK25 undergoes autophosphorylation in response to TNF-α and nerve growth factor (NGF), respectively (25, 35). While STK25 becomes activated in multiple cell types with a variety of stimuli, we have demonstrated that STK25 also undergoes autophosphorylation and activation in THP-1 cells in response to TLR4, TLR7/8, and TLR9 engagement. Furthermore, we detected an increase in STK25 protein expression at 24 h post-stimulation with R848 in THP-1 cells (Figure 3C-D). The induction of STK25 protein expression by R848 supports the notion of a positive feedback loop that drives STK25 activity in the context of TLR activation. However, the transcriptional mechanisms responsible for the upregulation of STK25 downstream of TLR7/8 have never been studied.

The canonical pathway of TLR-induced IRF5 activation involves phosphorylation, dimerization, and nuclear translocation (16, 33). Given the ability of STK25 to modulate IRF5 phosphorylation and transcriptional activation, we evaluated the role of STK25 in TLR-induced IRF5 nuclear translocation in murine primary immune cells. Importantly, we observed a reduction in R848- and CpG-B-induced IRF5 nuclear translocation in KO peripheral blood monocytes and B cells compared to WT (Figure 3F-G). In addition, we confirmed the importance of STK25 in TLR-mediated proinflammatory cytokine production through studies with KO splenocytes. Accordingly, the production of IL-6 downstream of R848 and CpG-B ligation was attenuated in KO splenocytes compared to WT (Figure 3J-K). These findings support our initial studies in human primary cells and highlight the ability of STK25 to regulate multiple steps involved in TLR-mediated IRF5 function, from upstream phosphorylation to downstream proinflammatory cytokine production.

Throughout our studies, the mechanisms by which STK25 engages with IRF5 downstream of TLR7/8 activation have remained elusive. The association between a kinase and its substrate can be difficult to capture, due to the transient nature of the phosphorylation process. However, the association between PYK2 and IRF5 prompted our investigation into whether STK25 binds to IRF5 (26). In immunoprecipitation studies in THP-1 cells and Ramos B cells, we were unable to observe an association between STK25 and IRF5. Given that the GCK-III subfamily members, MST3 and MST4, modulate signal transduction pathways through direct interactions with IKKβ and TRAF6, respectively, it is possible that STK25 is localized to IRF5 through an interaction with a TLR signaling component (36, 37). Alternately, the targeting of STK25 to IRF5 may depend on an unknown TLR adaptor protein. Accordingly, the recently identified adaptor protein, TASL, interacts with IRF5 and facilitates the recruitment of IKKβ downstream of TLR engagement (38).

Although activation of IRF5 in response to a microbial insult can aid in pathogen elimination, the dysregulation of IRF5 activity has been implicated in the pathogenesis of several autoimmune conditions. Specifically, *IRF5* genetic polymorphisms have been associated with aberrant leukocyte function in SLE through hyperactivation of IRF5 (20). Interestingly, the genetic ablation of *Irf5* has been found to confer protection in several murine models of SLE, including NZB/W F1, MRL/lpr, *Lyn*^*-/-*^, and pristane-induced mice (21, 22, 23). As such, IRF5 has emerged as a therapeutic target for the treatment of SLE and other autoimmune conditions. While the chemical inhibition of IRF5 remains an efficacious therapeutic strategy, the modulation of IRF5 kinases represents an important avenue for disrupting IRF5 function. The recent development of small molecule inhibitors of STK25 provides an invaluable research tool and represents an important step toward the production of clinically relevant therapeutic agents (39). Even though STK25 has never been directly implicated in the pathogenesis of SLE, the ability of STK25 to regulate the TLR-induced activation of IRF5 indicates that STK25 may contribute to the hyperactivated IRF5 phenotype in this disease. Quite strikingly, we observed a substantial increase in p-STK25 expression in SLE PBMCs compared to healthy controls at steady-state, suggesting that STK25 is basally activated in SLE (Figure 4A-C). These data support the further examination of STK25 function in models of murine lupus. In conclusion, the identification of STK25 as a positive regulator of IRF5 in SLE could lead to the development of targeted therapeutic agents that could selectively inhibit IRF5 hyperactivation to reduce inflammation and disease burden.

## Materials and Methods

### Animals

*Stk25*-deficient (*Stk25*^*-/-*^) mice were generated by Cre-mediated excision of exons 4 and 5 and genotyped as previously described (10). Deletion of *Stk25* was confirmed at the protein level by immunoblot analysis of *Stk25*^*+/+*^ and *Stk25*^*-/-*^ splenocytes (Figure 3E). All mice used in this study were between 8-12 weeks of age. All animal care and experimental procedures were conducted in accordance with the *Guide for the Care and Use of Laboratory Animals* (National Academies Press, 2011) and approved by the IACUC of the Feinstein Institutes for Medical Research.

### Cell culture

THP-1 cells (TIB-202), Ramos cells (CRL-1596), Jurkat cells (TIB-152), and HEK293T cells (CRL-3216) were obtained from ATCC. Cells were certified mycoplasma-free at the time of receipt from ATCC. THP-1, Ramos, and Jurkat cells were maintained in RPMI 1640 medium supplemented with 10% fetal bovine serum (FBS) and 1X penicillin/streptomycin (P/S). HEK293T cells were maintained in DMEM (high glucose, GlutaMAX) supplemented with 10% FBS and 1X P/S. Cells were incubated at 37°C in a 5% CO_2_ environment. For kinetic studies of p-STK25 (T174) expression, cells were seeded in 12-well plates at a density of 1 x 10^6^ cells per well in 1 mL of complete RPMI 1640 and stimulated with R848 (1 μg/mL), CpG-B (2.5 μg/mL), or LPS (1 μg/mL) accordingly. For STK25 expression studies, cells were seeded in 12-well plates at a density of 1 x 10^6^ cells per well in 1 mL of complete RPMI 1640 and stimulated with R848 (1 μg/mL) for 24 h.

### Kinome-wide siRNA screen

A kinome-wide siRNA library that contained 4 individually arrayed siRNA sequences in 384-well plates was purchased from Qiagen. The library consisted of known kinases and associated proteins. To screen the library, 5 μL of each siRNA at 200 nM in Opti-MEM (Gibco, 31985062) were added to 384-well plates followed by the addition of 5 μL of Lipofectamine 3000 RNAiMax Transfection Reagent (Invitrogen, 13778030), at 12.5 μL/mL. After 30 min of incubation at RT, THP-1 cells were seeded at a density of 20 x 10^4^ cells per well in 40 μL of RPMI 1640 supplemented with 1% FBS. After 48 h, the cells were stimulated with R848 (30 μM) for 24 h. Culture supernatants were harvested post-stimulation and the production of TNF-α was assessed using the AlphaLISA Immunoassay Kit (PerkinElmer, AL208C) according to the manufacturer’s protocol. Cell viability was assessed using the CellTiter-Glo Luminescent Viability Assay (Promega, G7570).

### Human primary monocyte isolation

Human primary monocytes were isolated from healthy donors using the autoMACS Pan Monocyte Isolation Kit (Miltenyi Biotec, 130-096-537). Briefly, one 40 mL Leukopak of blood was diluted with 100 mL of Dulbecco’s phosphate-buffered saline (DPBS) + 2% FBS + 1 mM EDTA. A 35 mL aliquot of the diluted sample was transferred to a 50 mL ACCUSPIN tube (Sigma-Aldrich, A2055) containing 15 mL of Histopaque-1077 (Sigma-Aldrich, 10771). After centrifugation, the interphase layer containing the peripheral blood mononuclear cells (PBMCs) was transferred to a clean 50 mL tube and washed twice with DPBS + 2% FBS + 1 mM EDTA. Monocytes were isolated from the PBMC fraction through negative selection by depleting the cells labeled with CD3, CD7, CD16, CD19, CD56, CD123 and Glycophorin A using a MACS Separator (Miltenyi Biotec).

### Generation of human primary monocyte-derived dendritic cells (MDDCs)

Human primary monocytes were seeded at a density of 5 x 10^5^ cells per well in 1 mL of RPMI 1640 supplemented with 25 ng/mL of IL-4 and 50 ng/mL of GM-CSF. The media was replaced every 2 to 3 days. MDDCs were harvested after 7 days of differentiation.

### Human primary cell transfection

The siRNA constructs (Qiagen) were mixed with 0.14 μL of Lipofectamine RNAiMAX Transfection Reagent (Invitrogen, 13778030) and incubated at RT for 20 to 30 min. The siRNA (20 nM)/lipofectamine(1.4 μl/ml) complex was then transferred to monocytes or MDDCs seeded at the density of 50,000/well and 25,000/well in 100 μl, respectively. The complex was then added to human primary monocytes or MDDCs that were seeded at a density of 5 x 10^4^ cells per well or 2.5 x 10^4^ cells per well in 100 μL of RPMI 1640, respectively. At 48 h post-transfection, the cells were stimulated with R848 (30 μM) for 24 h. Culture supernatants were harvested post-stimulation for analysis of proinflammatory cytokine production by AlphaLISA Immunoassay Kits.

### AlphaLISA immunoassay

The expression of IL-6 and TNF-α in culture supernatants was detected using the corresponding AlphaLISA Immunoassay Kit (PerkinElmer, AL223C and AL208C) according to the manufacturer’s protocol.

### Immunoblot analysis

Briefly, whole cell lysates were prepared by lysing cells in NP-40 lysis buffer (50 mM Tris-HCl (pH 7.4), 150 mM NaCl, 1% NP-40 and 5 mM EDTA) (Thermo Scientific, J60766-AP) supplemented with Halt Protease Inhibitor Cocktail (Thermo Scientific, 87786) and PhosStop Phosphatase Inhibitor Cocktail (Roche, 4906845001). Sample protein concentrations were quantified using the DC protein assay (Bio-Rad, 5000112). In general, 15-25 μg of protein per sample were separated by SDS-PAGE using the Bolt Bis-Tris system (Invitrogen). Proteins were transferred to 0.45 μm nitrocellulose membranes (MDI, SCNX8402XXXX101) using a wet tank transfer system. Transfer efficiency was assessed by incubating the membranes in 5 mL of Ponceau S (Sigma-Aldrich, P7170) for 5 min followed by destaining with Tris-buffered saline-0.05% Tween 20 (TBST) (Thermo Scientific, J77500.K2). The membranes were blocked for 1 h at RT with 5% bovine serum albumin (BSA) in TBST and incubated overnight at 4°C with the primary antibody diluted in the blocking buffer. The membranes were washed three times for 5 min each with TBST and incubated with the secondary antibody diluted in the blocking buffer for 1 h at RT. The membranes were washed three times for 5 min each with TBST and incubated with 1 mL of chemiluminescent detection reagent (Cytiva, RPN2232) for 3 min before image acquisition using a ChemiDoc MP Imaging System (Bio-Rad Laboratories). Horseradish peroxidase (HRP)-conjugated antibodies against β-actin (Cell Signaling, 12620, 1:5000) and GAPDH (Cell Signaling, 3683, 1:5000) were used as loading controls for protein normalization. Densitometric analysis was performed using the Image Lab software (Bio-Rad Laboratories).

### Immunoblot antibodies

*Target (Vendor, Catalog Number, Primary Dilution, Secondary Dilution, Buffer)*

IRF5 (Abcam, ab181553, 1:1000, 1:10,000, 5% BSA/TBST)

p-Threonine (Cell Signaling, 9386, 1:1000, 1:10,000, 5% BSA/TBST)

STK25 (Abcam, ab157188, 1:1000, 5% BSA/TBST)

p-STK25 (T174) (Abcam, ab76579, 1:1000, 5% BSA/TBST)

β-actin HRP conjugate (Cell Signaling, 12620, 1:5000, n/a, 5% BSA/TBST)

GAPDH HRP conjugate (Cell Signaling, 3683, 1:5000, n/a, 5% BSA/TBST)

Secondary: IgG HRP conjugate (Cell Signaling, 7074)

### Scintillation proximity assay (SPA)

Recombinant human IRF5 protein (residues 222-467; 3 μM) conjugated to biotin was incubated with recombinant human STK25, STK16, SPHK2, or MAP3K19 protein in the presence of isotopically-labeled ATP (γ-^33^P) for 1 h. Samples were incubated with streptavidin-conjugated scintillation beads and γ-^33^P incorporation was determined as described (40, 41).

### *In vitro* kinase assay

Recombinant human IRF5 protein (Abcam, ab173024) (500 ng) was incubated with recombinant human STK25 protein (Signalchem, S43-10G-10) (350 ng) in kinase reaction buffer (40 mM Tris-HCl pH 7.4, 20 mM MgCl_2_, 0.1 mg/ml BSA, 0.05 mM DTT, and 0.05 mM ATP) for 1 h at RT. The 10 μL reactions were quenched via the addition of 2X sample buffer, separated by SDS-PAGE, and transferred to nitrocellulose membranes. For the ADP-Glo kinase assays (Promega, TM313), 5 μL reactions were prepared with varying concentrations of recombinant human IRF5 protein (Abcam, ab173024) and recombinant human STK25 protein (Signalchem, S43-10G-10) or recombinant human IKKβ protein (Signalchem, I03-10BG-10) in kinase reaction buffer in 96-well plates. The reactions were incubated for 1 h at RT and quenched via the addition of 5 μL of ADP-Glo Reagent to deplete the unconsumed ATP. The 10 μL reactions were incubated for 40 min at RT followed by the addition of 10 μL of Kinase Detection Reagent to convert ADP to ATP. The 20 μL reactions were incubated for 30 min at RT and the luminescence of each reaction was measured with a plate-reading luminometer with an integration time of 1 sec per well.

### Phos-tag immunoblot analysis

The Phos-tag immunoblotting method was described previously (42). Briefly, quenched kinase reactions (20 μL) were subjected to Phos-tag SDS-PAGE using a 7.5% SuperSep Phos-tag (50 μm/L) for 30 min at 15 mA and then 100 min at 30 mA. Gels were incubated three times in 25 mL of EDTA-containing transfer buffer (48.2 mM Tris, 38.9 mM glycine, 0.037% SDS, 20% methanol, 10 mM EDTA) for 20 min each with agitation. After washing the gels once for 20 min in EDTA-free transfer buffer (48.2 mM Tris, 38.9 mM glycine, 0.037% SDS, 20% methanol), proteins were transferred to 0.45 μm nitrocellulose membranes (MDI, SCNX8402XXXX101) using a wet tank transfer system. Transfer efficiency was assessed by incubating the membranes in 5 mL of Ponceau S (Sigma-Aldrich, P7170) for 5 min followed by destaining with Tris-buffered saline-0.05% Tween 20 (TBST) (Thermo Scientific, J77500.K2). The membranes were blocked for 1 h at RT with 5% bovine serum albumin (BSA) in TBST and incubated overnight at 4°C with the anti-IRF5 antibody diluted 1:1000 in the blocking buffer. The membranes were washed three times for 5 min each with TBST and incubated with anti-Rabbit IgG secondary antibody diluted 1:10,000 in the blocking buffer for 1 h at RT. The membranes were washed three times for 5 min each with TBST and incubated with 1 mL of chemiluminescent detection reagent (Cytiva, RPN2232) for 3 min before image acquisition using a ChemiDoc MP Imaging System (Bio-Rad Laboratories).

### Mass spectrometry

Kinase reactions (10 μL) were briefly subjected to SDS-PAGE for 5 min using the Bolt Bis-Tris system (Invitrogen). The gel was stained with 25 mL of InstantBlue Coomassie Protein Stain (Abcam, ab119211) for 15 min at RT and protein bands of interest were excised. Gel slices were incubated with 1 mL of 30% ethanol for 20 min at 70°C. After three washes with 30% ethanol, the gel slices were stored in 1 ml of deionized H_2_O at 4°C until downstream analysis. Gel slices were washed three times with 100 μL of 50 mM ABC/25% ACN (100 mM ammonium bicarbonate/25% acetonitrile) for 5 min each at 55°C with agitation. Gel slices were dehydrated with 100 μL of ACN. ACN was removed and the gel slices were dehydrated by vacuum centrifugation for 10 min. Gel slices were reconstituted in 50 μL of fresh 3 mM Tris(2-carboxyethyl)phosphine (TCEP)/50 mM ABC and incubated at 55°C for 20 min. Excess TCEP solution was removed and 50 μL of fresh 10 mM CEMTS/EtOH was added to the gel slices and subsequently incubated at 55°C for 20 min. Excess CEMTS solution was removed and the gel slices were washed three times with 100 μL of 50 mM ABC/25% ACN for 10 min at 55°C with agitation. Gel slices were dehydrated with 100 μL of ACN. ACN was removed and gel slices were dehydrated by vacuum centrifugation for 10 min. Gel slices were incubated with 5 μL of sequencing grade modified procine trypsin (500 ng) in 50 mM ABC for 5 min on ice. Gel slices were incubated with 30 μL of 0.02% ProteaseMAX Surfactant in 50 mM ABC for 10 min on ice. Proteins were digested at 37°C overnight. Gel slices were incubated with 50 μL of 80% ACN/1% TFA for 15 min at 55°C with agitation. Samples were subjected to two elution steps with subsequent pooling and lyophilization by vacuum centrifugation. Peptide pellets were resuspended in 20 μL of Loading Buffer (5% DMSO, 0.1% TFA in water) and 3 μL of each sample was injected. Peptides were loaded on a 30 cm x 75 μm ID column packed with Reprosil 1.9 C18 silica particles (Dr. Maischt), and resolved on a 8-35% acetonitrile gradient in water (0.1% formate) at 250 nl/min. Eluting peptides were ionized by electrospray (2200V) and transferred into an Exploris 480 mass spectrometer (Thermo). Exploris 480 MS fully recalibrated, mass error <2ppm. In nano-spray mode, >2500 proteins and 15000 peptides identified (FDR<1%) from 200ng HeLa tryptic proteome. Nanoscale RP: <45 s peak width, >60 minutes effective elution during 90 minutes gradient. The water blank demonstrated no evident carry over. The MS was set to collect 120K resolution precursor scans (m/z 380-2000 Th) and 60K HCD fragmentation spectra at stepped 28,33,38% normalized energy, with first mass locked to 100 Th. Files searched using the Mascot scoring function within ProteomeDiscoverer, with mass tolerance set at 5ppm for MS1, and 0.01 Da for MS2. Spectra were matched against the UniProt human consensus databases plus a database of common contaminants (cRAP). M-oxidation, N/Q-deamidation, and S/T/Y-phosphorylation were set as variable modifications. Peptide-spectral matches were filtered to maintain FDR<1%.

### Reporter assay

HEK293T cells were seeded in white, tissue culture-treated 96-well plates at a density of 2.5 x 10^4^ cells per well in 100 μL of DMEM and incubated at 37°C for 24 h. Cells were co-transfected with 100 ng of each target plasmid along with 100 ng of pGL3-ISRE-Luc and 20 ng of pRL-CMV using Lipofectamine 3000 Transfection Reagent (Invitrogen, L3000015). Cells were lysed 24 h post-transfection using the Dual-Glo Luciferase Assay System (Promega, E2920) and luminescence was analyzed using a plate reader. The firefly luciferase signal was normalized to the *Renilla* luciferase signal to determine the relative luciferase units (RLU).

**Table 1.**
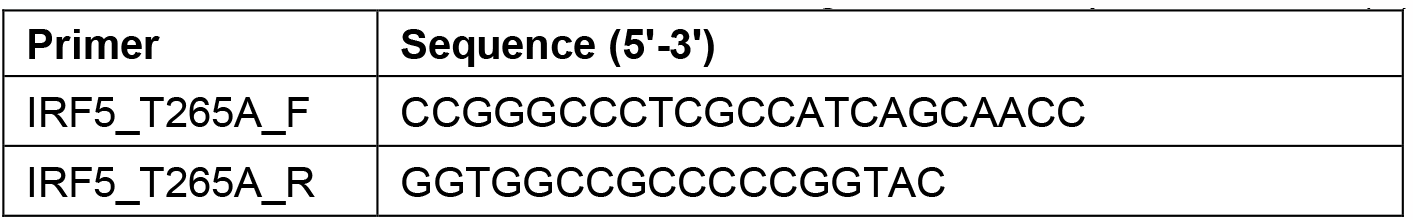
Primers used for site-directed mutagenesis of the pcDNA3.1+/C-(K)-DYK-IRF5 vector.

### Plasmids

The cDNA of human IRF5 isoform b (RefSeq accession no. NM_032643.4) was cloned into the pcDNA3.1+/C-(K)-DYK vector with a C-terminal FLAG tag (GenScript). The cDNA of human STK25 isoform 1 (RefSeq accession no. NM_001271977.2) was cloned into the pcDNA3.1+ vector (GenScript). The Q5 Site-Directed Mutagenesis Kit (NEB) was used to generate pcDNA3.1+/C-(K)-DYK-IRF5-T265A. The pGL3-ISRE-Luc vector and the pRL-CMV vectors were obtained from Promega.

### Multiple sequence alignment

Protein sequences were extracted from the UniProt database and aligned with ClustalOWS and rendered in Jalview (41).

### Flow cytometry Spleen

Spleens from WT and *Stk25*^*-/-*^ mice were harvested and homogenized with frosted slides in 4 mL of DPBS + 2% FBS. The samples were incubated with 5 mL of 1X RBC Lysis Buffer for 5 min on ice. Samples were washed with 5 mL of Dulbecco’s PBS (DPBS) + 2% FBS and pelleted by centrifugation. Pellets were resuspended in 10 mL of DPBS + 2% FBS and filtered through a 70 μm cell strainer. The total number of cells in each sample was determined using a hemocytometer and adjusted to a final concentration of 4 x 10^6^ cells/mL in complete RPMI 1640 medium. Cells were plated in an ultralow adherent 24-well plate and stimulated with R848 (1 μg/mL), CpG-B (2.5 μg/mL), or LPS (100 nM/mL) for 18 h. Cells were incubated with Brefeldin A for 2 h to facilitate the intracellular accumulation of IL-6. Cells were harvested, stained for surface markers in the dark, and subjected to fixation and permeabilization with 0.1% Triton X-100 in 2% formaldehyde/PBS for 20 min. Cells were then stained for intracellular IL-6 and events were collected using the BD LSR Fortessa cell analyzer (BD Biosciences). Data was analyzed using the FlowJo Software (BD Biosciences).

### Antibodies

*Target-Conjugate (Vendor, Catalog Number, Clone, Volume)*

CD45-APC/Cy7 (BioLegend, 103116, 30-F11, 0.25 μL)

CD11b-PE (BioLegend, 101208, M1/70, 0.25 μL)

B220-BV510 (BioLegend, 103248, RA3-6B2, 0.25 μL)

Ly6G-PerCP/Cy5.5 (BioLegend, 127615, 1A8, 0.25 μL)

Ly6C-PE/Cy7 (BioLegend, 128018, HK1.4, 0.25 μL)

CD4-BV421 (BioLegend, 100438, GK1.5, 0.25 μL)

IL-6-APC (BioLegend, 504508, MP5-20F3, 1 μL)

TNF-α-FITC (BioLegend, 506304, MP6-XT22, 1 μL)

### Imaging flow cytometry

#### Peripheral blood cells

Imaging flow cytometry was performed as previously described on the Amnis ImageStream (16). Briefly, peripheral blood was isolated from WT and *Stk25*^*-/-*^ mice via cardiac puncture and incubated with 5 mL of 1X RBC Lysis Buffer for 5 min on ice. Samples were washed with 5 mL of PBS, and the total number of cells in each sample was determined using a hemocytometer. Samples were adjusted to a final concentration of 0.5 x 10^6^ cells/mL in complete RPMI 1640 medium. Cells were plated in a 24-well plate and stimulated with R848 (1 μg/mL) or CpG-B (2.5 μg/mL) for 2 h. Samples were washed with 5 mL of PBS, resuspended in 100 μL of PBS + 2% bovine serum albumin (BSA), and stained for surface markers for 40 min at 4°C in the dark. Samples were washed with 5 mL of PBS and pellets were resuspended in 200 μL of 4% paraformaldehyde (PFA)/PBS. Samples were incubated for 1 h at RT and washed twice with 5 mL of PBS. For the permeabilization step, samples were resuspended in 300 μL of 0.5% Triton X-100 (200 μL) + 2 μL of Fc Shield (anti-mouse CD16/CD32, Tonbo Biosciences, 70-0161-U500) + 100 μL of 5% BSA/PBS and incubated overnight at 4°C in the dark. The next morning, samples were washed with 5 mL of 0.1% Triton X-100/PBS and incubated with 1 μL of IRF5 antibody (Abcam, 181553) in 100 μL of 0.1% Triton X-100/PBS for 2 h at RT. Samples were washed three times with 5 mL of 0.1% Triton X-100/PBS and incubated with 2 μL of anti-rabbit IgG secondary-APC (Invitrogen, A-10931) in 100 μL of 0.1% Triton X-100/PBS for 40 min at RT. Samples were washed twice with 5 mL of 0.1% Triton X-100/PBS and nuclei were stained with DAPI (Invitrogen, D1306) for 5-8 min at RT. Samples were washed once with 10 mL of PBS, resuspended in 50 μL of 4% PFA/PBS, and transferred to 1.5 mL microcentrifuge tubes. Events were collected using the Cytek Amnis ImageStream Mk II imaging flow cytometer (Cytek Biosciences). Data was analyzed using the IDEAS Image Analysis Software (Cytek Biosciences).

### Antibodies

*Target-Conjugate (Vendor, Catalog Number, Clone, Volume)*

B220-FITC (BD Pharmingen, 553088, RA3-6B2, 2 μL) CD4-BV510 (BioLegend, 100559, RM4-5, 0.7 μL)

CD8a-APC/Cy7 (BioLegend, 100714, 53-6.7, 1 μL)

CD11b-PE (BioLegend, 101208, M1/70, 1 μL)

CD11c-PE/Cy7 (BioLegend, 117318, N418, 1 μL)

Anti-rabbit IgG-APC (Invitrogen, A-10931, Polyclonal, 2 μL)

### ELISA

Splenocytes from WT and *Stk25*^*-/-*^ mice were plated at a density of 4 x 10^6^ cells/mL in 24-well plates and stimulated with R848 (1 μg/mL), CpG-B (2.5 μg/mL), or LPS (100 nM/mL) for 24 h. The expression of IL-6 in culture supernatants was detected using the Murine IL-6 Mini ABTS ELISA Development Kit (PeproTech, 900-M50) according to the manufacturer’s protocol.

### Statistical analysis

The statistical measure of SSMD was used to evaluate positive hits from the siRNA screen for confirmatory studies. For the primary screen in THP-1 cells, the robust version of SSMD was used on a plate-by-plate basis using the siRNA sequences on that plate as the negative reference. For the confirmatory screen in human primary cells, additional negative control siRNAs were included as the negative reference. A two-tailed Student’s t test was used for comparisons between two samples with normal distribution. A one-way ANOVA was used to compare one factor across multiple groups. Afterwards, Tukey’s multiple comparison test was used to detect differences between groups. GraphPad Prism 9 (GraphPad Software) was used for statistical analysis and figure generation. A P value of < 0.05 was considered statistically significant.

## Acknowledgments

The authors would like to thank Dr. Seng-Lai (Thomas) Tan and Dr. Gang Chen for helpful discussions and sharing of the library screening data. The authors would also like to thank Dr. Paolo Cifani and Dr. Darryl Pappin of the Mass Spectrometry Shared Resource at Cold Spring Harbor Laboratory for their efforts on the phosphopeptide mapping studies.

