## Supplemental Data for "STK25 is an IRF5 kinase that promotes TLR7/8-mediated inflammation"

### Supporting Information Figure Legends

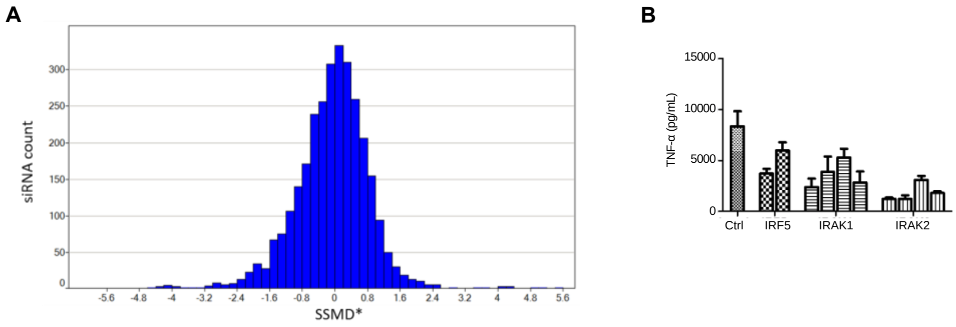


**Figure S1. Validation of the kinome-wide siRNA screen in THP-1 cells.**
**A**, Normal distribution of strictly standardized mean difference (SSMD) scores for each siRNA construct. An SSMD score of -1 for a particular siRNA molecule was considered a positive hit and targets with 2 or more hits were selected for validation in follow up studies. **B**, Production of TNF-α by THP-1 cells stimulated with R848 for 24 h following siRNA knockdown of *IRF5*, *IRAK1*, or *IRAK2* (*N* = 3 technical replicates with 2-4 distinct siRNA constructs per target). Data represent mean ± SEM.

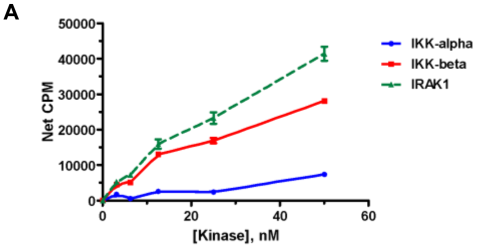


**Figure S2. IRAK1 and IKKβ phosphorylate IRF5 in vitro. A**, Phosphorylation of a biotinylated C-terminal construct of IRF5 (residues 222-467) by kinases involved in TLR signaling in an in vitro scintillation assay (*N* = 4 independent replicates). Data represent mean ± SEM.

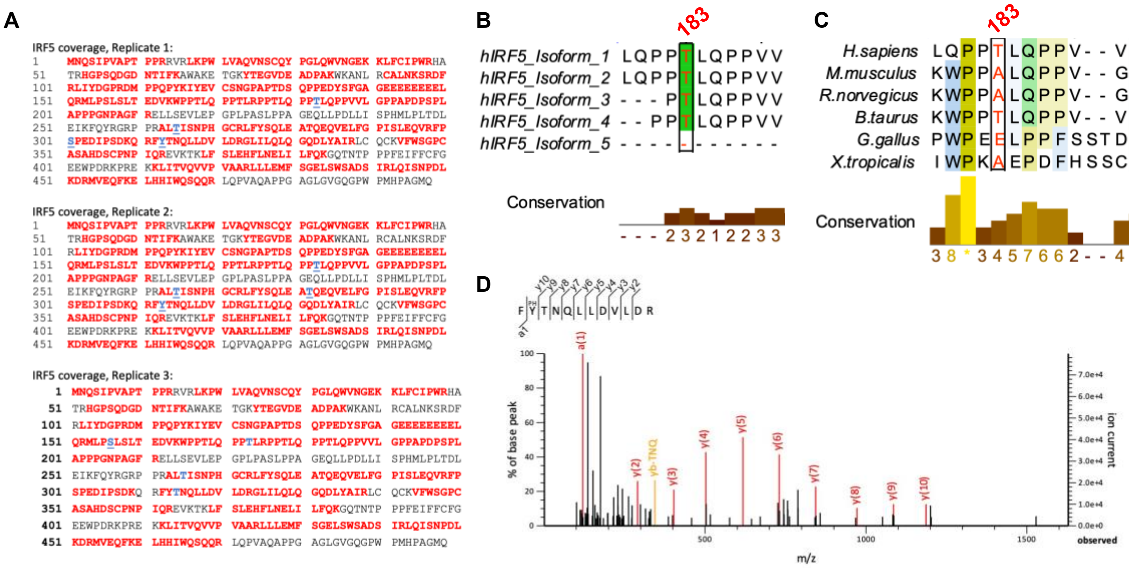


**Figure S3. Identification of STK25-mediated IRF5 phosphorylation sites by mass spectrometry. A**, IRF5 peptide coverage from three independent replicates. Residues in red were detected and residues in blue were determined to be phosphorylated. **B**, Conservation of Thr183 across multiple human isoforms of IRF5. **C**, Conservation of Thr183 in IRF5 protein sequences from multiple species. **D**, Spectral data for the phosphorylation of IRF5 at Tyr313.

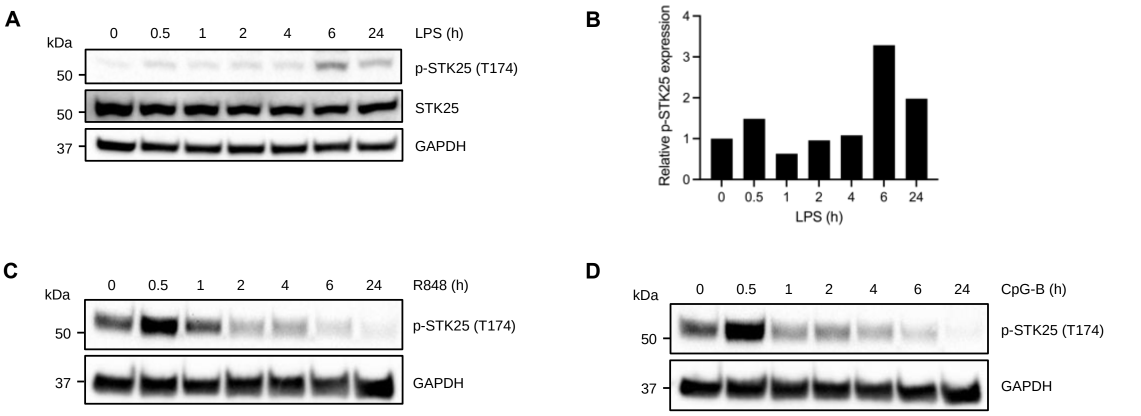


**Figure S4. STK25 undergoes autophosphorylation in response to TLR4 activation in THP-1 cells. A**, Immunoblot analysis of STK25 autophosphorylation at Thr174 in THP-1 cells stimulated with LPS for 0.5, 1, 2, 4, 6, or 24 h. Representative of 2 independent experiments. Blots were probed with antibodies against phospho-STK25 (T174), STK25, and GAPDH. **B**, Representative densitometric analysis of p-STK25 (T174) protein levels after normalization to total STK25 protein levels and the expression of GAPDH. **C**, Immunoblot analysis of STK25 autophosphorylation at Thr174 in Ramos B cells stimulated with R848 for 0.5, 1, 2, 4, 6, or 24 h. Representative of 2 independent experiments. Blots were probed with antibodies against phospho-STK25 (T174) and GAPDH. **D**, Immunoblot analysis of STK25 autophosphorylation at Thr174 in Ramos B cells stimulated with CpG-B for 0.5, 1, 2, 4, 6, or 24 h. Representative of 2 independent experiments. Blots were probed with antibodies against phospho-STK25 (T174) and GAPDH.
